# Localization of Proteins Involved in the Biogenesis of the Photosynthetic Apparatus to Thylakoid Subdomains in Arabidopsis

**DOI:** 10.1101/2024.06.21.600055

**Authors:** Prakitchai Chotewutmontri, Alice Barkan

**Affiliations:** Institute of Molecular Biology, University of Oregon, Eugene, OR 97403

**Author notes:** Crop Improvement and Genetics Research, Western Regional Research Center, United States Department of Agriculture—Agricultural Research Service, Albany, CA 94710.

**Keywords:** thylakoid membrane, thylakoid margins, thylakoid curvature, chloroplast, photosystem biogenesis, photosystem II repair

## Abstract

Thylakoid membranes in chloroplasts and cyanobacteria harbor the multisubunit protein complexes that catalyze the light reactions of photosynthesis. In plant chloroplasts, the thylakoid membrane system comprises a highly organized network with several subcompartments that differ in composition and morphology: grana stacks, unstacked stromal lamellae, and grana margins at the interface between stacked and unstacked regions. The localization of components of the photosynthetic apparatus among these subcompartments has been well characterized. However, less is known about the localization of proteins involved in the biogenesis and repair of the photosynthetic apparatus, the partitioning of proteins between two recently resolved components of the traditional margin fraction (refined margins and curvature), and the effects of light on these features. In this study, we analyzed the partitioning of numerous thylakoid biogenesis and repair factors among grana, curvature, refined margin, and stromal lamellae fractions of Arabidopsis thylakoid membranes, comparing the results from illuminated and dark-adapted plants. Several proteins previously shown to localize to a margin fraction partitioned in varying ways among the resolved curvature and refined margin fractions. For example, the ALB3 insertase and FtsH protease involved in photosystem II (PSII) repair were concentrated in the refined margin fraction, whereas TAT translocon subunits and proteins involved in early steps in photosystem assembly were concentrated in the curvature fraction. By contrast, two photosystem assembly factors that facilitate late assembly steps were depleted from the curvature fraction. The enrichment of the PSII subunit OE23/PsbP in the curvature fraction set it apart from other PSII subunits, supporting the previous conjecture that OE23/PsbP assists in PSII biogenesis and/or repair. The PSII assembly factor PAM68 partitioned differently among thylakoid fractions from dark-adapted plants and illuminated plants, and was the only analyzed protein to convincingly do so. These results demonstrate an unanticipated spatial heterogeneity of photosystem biogenesis and repair functions in thylakoid membranes, and reveal the curvature fraction to be a focal point of early photosystem biogenesis.

## Introduction

The light reactions of oxygenic photosynthesis take place in thylakoid membranes of cyanobacteria and chloroplasts. Thylakoid membrane networks in plant chloroplasts include two major morphological features: tightly appressed stacked disks called grana and unstacked membranes known as stromal lamellae. Stromal lamellae connect with grana through slit-like "margins", such that the network encloses a single continuous lumenal space (Anderson and Andersson, 1988; Austin and Staehelin, 2011; Daum and Kuhlbrandt, 2011; Ruban and Johnson, 2015; Kirchhoff, 2019; Rantala et al., 2020). The multisubunit complexes that catalyze photosynthetic electron transport and ATP synthesis are heterogeneously distributed within this network (Anderson and Andersson, 1988; Koochak et al., 2019; Trotta et al., 2019; Höhner et al., 2020): Photosystem II (PSII) is predominantly located in the grana, Photosystem I (PSI) and the ATP synthase are primarily found in stromal lamellae, while the cytochrome *b_6_f* complex (cyt *b_6_f*) exhibits a relatively uniform distribution. The partitioning of membranes between the appressed and nonappressed fractions as well as the dimensions of grana change in response to light conditions (Yoshioka-Nishimura et al., 2014; Yoshioka-Nishimura, 2016; Kirchhoff et al., 2017; Kirchhoff, 2019; Li et al., 2020; Rantala et al., 2020). These changes are believed to facilitate energy transduction, minimize PSII photodamage, and aid in the repair process should photodamage occur. For example, light-induced lumenal swelling may aid the diffusion of the electron carrier plastocyanin, while light-induced "fraying" of grana edges may enhance access of damaged PSII complexes to factors involved in PSII repair (Kirchhoff et al., 2011; Yoshioka-Nishimura, 2016; Höhner et al., 2020).

The proteins that make up the photosynthetic apparatus dominate the thylakoid proteome, but thylakoid membranes harbor hundreds of additional proteins that play vital roles in the targeting, assembly, protection, repair, and regulation of proteins involved directly in photosynthesis (Peltier et al., 2004; Flannery et al., 2021). Little is known about how the assembly and repair processes are orchestrated in three dimensional space. Proteins involved in these processes must have access to the chloroplast stroma, where plastid-encoded thylakoid proteins are synthesized and through which nucleus-encoded thylakoid proteins must transit. Indeed, a proteome analysis of separated grana and stromal lamellae showed biogenesis and repair factors to be concentrated in the latter fraction (Tomizioli et al., 2014). The interface between the grana and stromal lamellae is of particular interest because electrons must be shuttled between PSII in grana and PSI in stromal lamellae, and because mature PSII resides primarily in grana but its assembly and repair require exposure to the stroma. Photodamage to the PSII reaction center protein D1 triggers partial disassembly of PSII, proteolytic removal of damaged D1, its replacement with newly synthesized D1, and reassembly of a functional complex (reviewed in Jarvi et al., 2015; Theis and Schroda, 2016). The CP43-free PSII core that is produced following PSII photodamage, the FtsH protease that degrades photodamaged D1, and several PSII assembly/repair factors are enriched in thylakoid margins (Sakamoto et al., 2003; Puthiyaveetil et al., 2014; Yoshioka-Nishimura et al., 2014; Garcia-Cerdan et al., 2019; Koochak et al., 2019), suggesting the margins to be the site of PSII disassembly and reassembly during PSII repair.

In a recent advance, modifications to the methods used to purify grana margins resolved the traditional "margin" compartment into two components: a "curvature" domain encompassing the highly curved periphery of grana disks and a "refined margin" domain that connects grana to stromal lamellae (Koochak et al., 2019; Trotta et al., 2019; Rantala et al., 2020). CURT1 proteins, which help to establish the curvature at the periphery of grana disks, localize specifically to the curvature domain (Armbruster et al., 2013; Trotta et al., 2019), as does FZL, which is likewise implicated in the special architecture of this subdomain (Ogawa et al., 2023). However, very little information is available about the partitioning of photosystem biogenesis and repair factors between the curvature and refined margin fractions. Clarification of the spatial relationship among such proteins may elucidate assembly pathways and partnerships among proteins involved in these processes. To address this gap, we cataloged the partitioning of a wide array of proteins involved in the biogenesis and repair of the photosynthetic apparatus among grana, stromal lamellae, curvature, and refined margin fractions of Arabidopsis thylakoid membranes. Our survey included proteins involved in the targeting of proteins to the thylakoid membrane, the VAR1/FtsH protease involved in degradation of photodamaged D1 (Nishimura et al., 2016), ribosomal proteins, assembly factors for PSI and PSII, and more. We compared the partitioning of these proteins in thylakoid membranes isolated from dark-adapted and illuminated plants, motivated by evidence that light-induced unstacking of grana edges facilitates PSII repair (Puthiyaveetil et al., 2014). Our results show that proteins previously shown to copurify in the traditional margin fraction exhibit varying distributions among the refined margin and curvature fractions. With the notable exception of the PSII assembly factor PAM68, protein distributions did not change convincingly upon illumination. These fractionation behaviors provide insight into the spatial organization of photosystem biogenesis and repair, and reveal the curvature fraction to be of special relevance for early photosystem biogenesis.

## Materials and Methods

### Preparation of thylakoid membrane fractions

*Arabidopsis thaliana* Col-0 seedlings were germinated and grown on Murashige and Skoog (MS) medium containing 3% (w/v) sucrose and 3% (w/v) Phytagel under cycles of 10 h light (100 µmol photons m^-2^ s^-1^) and 14 h dark at 22°C for 14 days. Aerial tissues were harvested just before the end of the dark cycle (dark-adapted plants) or 3 h after dawn (illuminated plants). Dark-adapted plants were harvested under dim illumination (<1 µmol photons m^-2^ s^-1^) from a green headlamp and the tissues were kept in darkness when possible. Homogenization and fractionation of freshly harvested tissue was performed under the same dim light for dark-adapted tissue or under 100 µmol photons m^-2^ s^-1^ for illuminated tissue. Thylakoid subfractions were prepared as described by (Trotta et al., 2019) except that glyco-diosgenin (GDN) was used instead of digitonin.

Plant tissue was ground in ice-cold grinding buffer (50 mM HEPES-KOH pH 7.5, 330 mM sorbitol, 5 mM MgCl_2_, 2.5 mM ascorbate, 0.05% (w/v) BSA, 10 mM NaF), filtered through two sheets of Miracloth, and then centrifuged at 5,000 x g at 4°C for 4 min. The pellet was resuspended in osmotic shock buffer (50 mM HEPES-KOH pH 7.5, 5 mM sorbitol, 5 mM MgCl_2_, 10 mM NaF) and incubated on ice for 10 min before centrifugation at 5,000 x g at 4°C for 4 min. The pellet (total thylakoid fraction) was resuspended in storage buffer (50 mM HEPES-KOH pH 7.5, 100 mM sorbitol, 5 mM MgCl_2_, 10 mM NaF) and centrifuged at 5,000 x g at 4°C for 4 min. The thylakoid pellet was resuspended in GDN buffer (0.4% (w/v) GDN, 15 mM Tricine-KOH pH 7.8, 100 mM sorbitol, 10 mM NaCl, 5 mM MgCl2, 10 mM NaF) to reach a final chlorophyll concentration of 0.5 mg/mL, and solubilized for 8 min at room temperature before removal of unsolubilized material by centrifugation at 1,000 x g at 4°C for 3 min. Thylakoid subfractions were then collected by differential centrifugation: 10,000 x g for 30 min to pellet the grana core, followed by centrifugation of the supernatant at 40,000 x g for 30 min to pellet grana margins, and centrifugation of that supernatant at 144,000 x g for 1 h to collect the curvature fraction (loose pellet) and the stromal lamellae (pellet). Pellets were resuspended in GDN buffer and stored at -80°C. Protein concentrations were determined using the Pierce BCA Protein Assay (Thermo Fisher Scientific) and chlorophyll concentrations were determined with the method of (Porra et al., 1989).

### Immunoblotting and Antibodies

Proteins were separated by SDS-PAGE and analyzed by immunoblotting as described previously (Barkan, 1998) except for the use of precast Tris-glycine 4-20% polyacrylamide gels (Novex, Invitrogen). Antibodies to AtpB, D1, HCF106/TatB, HCF173, HCF244, OE23, OE33, PsaD, PetD, THA4/TatA were raised by our group and have been described previously (Walker et al., 1999; McDermott et al., 2019; Chotewutmontri et al., 2020). Antibody to OHP2 was raised in rabbits against a recombinant protein fragment corresponding to amino acids 39 through 135 of Arabidopsis OHP2 (gene model At1g34000.1). The crude OHP2 serum was affinity purified with the same antigen before use. Antibodies to PSA2 and PSA3 were described in (Fristedt et al., 2014) and (Shen et al., 2017), respectively, and are available from Agrisera under catalog numbers AS13 2654 and AS15 2872, respectively. Antibodies to ALB3 and cpFtsY were described in (Moore et al., 2000) and were generously provided by Misty Moore and Ralph Henry (University of Arkansas). Antibody to HCF136 was described in (Meurer et al., 1998) and was generously provided by Peter Westhoff (Heinrich Heine University, Düsseldorf). The cpSecY antibody was described in (Mori et al., 1999) and was generously provided by Carole Dabney-Smith (Miami University). Antibody to VAR1/FtsH5 was described in (Sakamoto et al., 2003) and was generously provided by Wataru Sakamoto (Okayama University). This antibody cross-reacts with FtsH1 but not with the co-paralogs VAR2 and FtsH8 (Sakamoto et al., 2003). The antibody to RBD1 was described in (Calderon et al., 2013) and was generously provided by Kris Niyogi (University of California). The antibody to TROL2 was generously provided by Yukari Asakura (formerly of Osaka University). Other antibodies were purchased from Agrisera [ChlG (AS14 2793), CURT1A (AS08 316), D2 (AS04 038), Lhca1 (AS01 005), Lhcb2 (AS01 003), PsaB (AS10 695), RPS1 (AS15 2875), RPS5 (AS15 3075), THF1 (AS07 240), Ycf3 (AS07 273)],or PhytoAB [PAM68 (PHY2289A)].

Gel loading of thylakoid subfractions was based on equal protein. The amount of chlorophyll (chl *a* +*b*) in each gel sample was as follows: total thylakoids, 1.4 µg light, 1.4 µg dark-adapted; refined margins, 0.7 µg light, 0.5 µg dark-adapted; granal core, 6.0 µg light, 8.3 µg dark adapted; curvature, 0.3 µg light, 0.2 µg dark adapted; stromal lamellae, 2.0 µg light, 1.3 µg dark-adapted.

## Results

### General properties of thylakoid subfractions

Thylakoid membranes were purified from the aerial portion of 2-week old Arabidopsis seedlings grown in diurnal cycles. Plants were harvested either 3 hours after illumination at dawn or shortly before the end of the night cycle (14 h in the dark). The membranes were fractionated as described by Trotta et al (Trotta et al., 2019), except that 0.4% (w/v) glyco-diosgenin (GDN) was used instead of 0.4% (w/v) digitonin. GDN is structurally similar to digitonin and has been shown to behave similarly to digitonin for the fractionation of thylakoid membranes into stromal lamellae, margins, and grana (Nishioka et al., 2021; Lee et al., 2022). Trotta and coworkers designated the material that pelleted during the initial low speed centrifugation (10,000 x g, 30 min) as grana core, the material that pelleted during subsequent centrifugation at 40,000g for 30 min as grana margins, and the material that pelleted during subsequent centrifugation at 144,000g for 1 h as stromal lamellae and curvature (tight and loose pellet, respectively). We follow that nomenclature here except that we use the term "refined margins" to distinguish this from the "margin" fraction recovered using prior fractionation protocols (e.g.Koochak et al., 2019; Nishioka et al., 2021), which includes the "curvature" domain (see (Trotta et al., 2019) and evidence below).

The chlorophyll *a/b* ratio was lowest in the grana core fraction (2.7) (Figure 1A), as expected due to enrichment of LHCII in grana. However, this ratio was somewhat higher than is typical (∼2.4) (Koochak et al., 2019; Trotta et al., 2019), suggesting that our grana core fraction was contaminated with nonappressed membranes. Other fractions were enriched in chlorophyll *a,* albeit with ratios that differed somewhat from the corresponding fractions reported previously (Trotta et al., 2019) (Figure 1A). Notably, the curvature fraction had a much higher chlorophyll *a/b* ratio than the refined margin fraction (Figure 1A) and a much higher ratio of protein to chlorophyll than any other fraction (Figure 1B). The chlorophyll *a/b* ratio of the curvature and stromal lamellae fractions decreased after illumination, as was observed previously for a traditional "margin" fraction that included the curvature domain (Trotta et al., 2019). Additionally, the protein-to-chlorophyll ratio in the curvature fraction increased upon illumination. These dynamic features of the curvature fraction correlate with changes in the morphology of grana edges in response to light (Kirchhoff, 2019).

**Figure 1.**
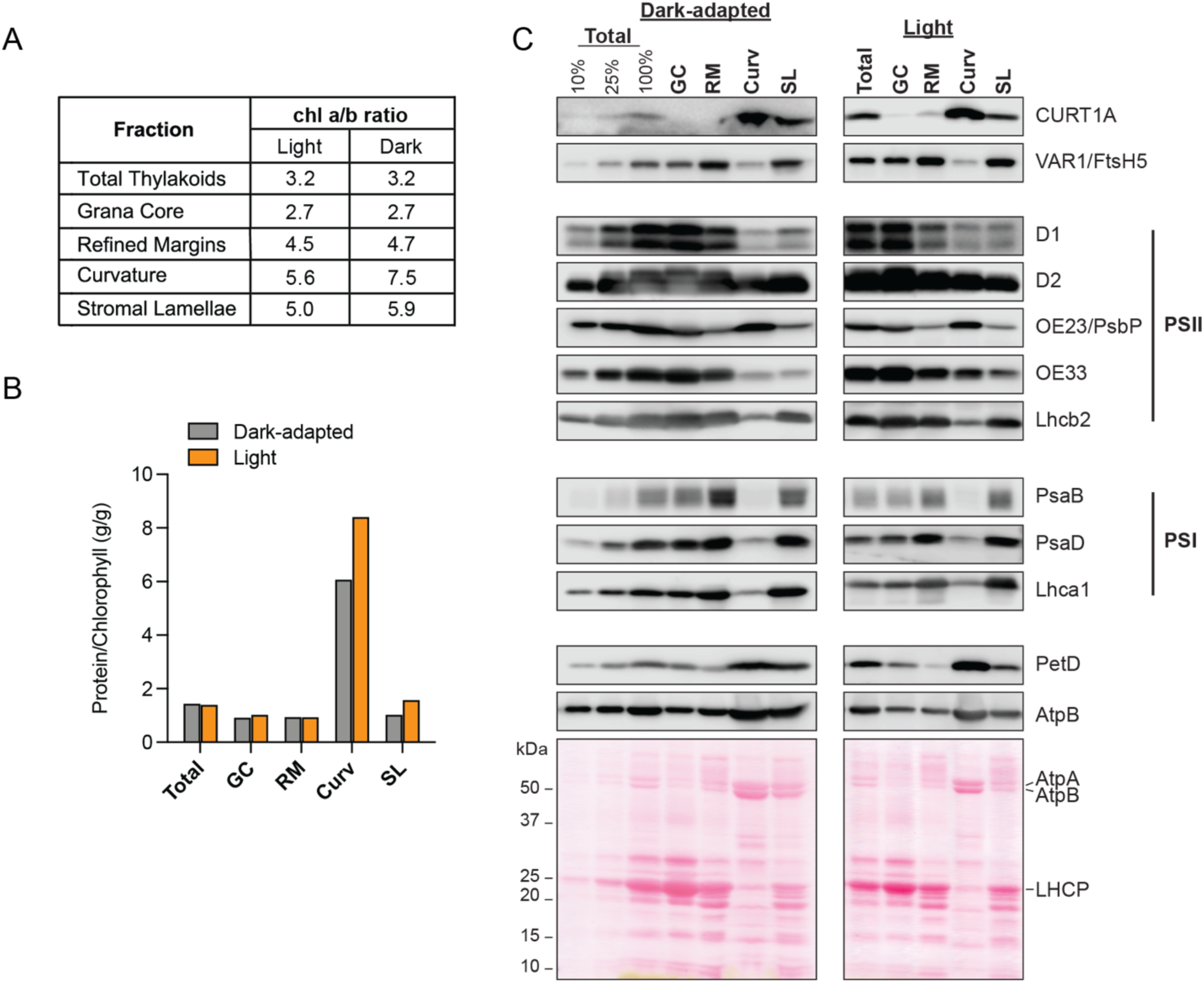
Characterization of thylakoid subfractions. (A) Chlorophyll *a/b* ratio of thylakoid membrane fractions. (B) Ratio of protein to chlorophyll in thylakoid membrane fractions. Total, total thylakoids; GC, grana core; RM, refined margins; Curv, curvature; SL, stromal lamellae. (C) Immunoblot analysis of markers for thylakoid subfractions. Lanes contained equal amounts of protein or the indicated dilutions of the total thylakoid sample. The amount of chlorophyll in each gel sample is provided in Materials and Methods. Replicate blots were probed with antibodies to the indicated proteins. D1 and D2 are subunits of the PSII reaction center. CURT1A is a marker for the curvature fraction. VAR1/Ftsh5 is a subunit of the thylakoid FtsH protease. OE23/PsbP and OE33 are subunits of the oxygen evolving complex associated with PSII. Lhcb2 is a subunit of the PSII light harvesting complex. PsaB and PsaD are reaction center subunits of PSI. Lhca1 is a subunit of the PSI light harvesting complex, PetD is a subunit of the cyt *b_6_f* complex. AtpB is a subunit of the thylakoid ATP synthase. Representative blots stained with Ponceau S are shown below, with prominent bands corresponding to subunits of the ATP synthase (AtpA, AtpB) and LHCII (LHCP) marked. Total, total thylakoids; GC, grana core; RM, refined margins; Curv, curvature; SL, stromal lamellae.

We further evaluated the composition of our fractions by immunoblot analysis of marker proteins whose localization to thylakoid subdomains had been studied previously (Figure 1C). The apparent partitioning of proteins among the fractions will differ depending upon how gel loading is normalized. It would be ideal to analyze the same proportion of each fraction, but this was impractical due to unknown losses from each fraction during the preparation. Gel loading based on equal chlorophyll inflates signals for fractions with a low representation of chlorophyll-rich complexes, whereas gel loading based on equal protein inflates signals from fractions with lower total protein content. Recognizing these limitations, we chose to normalize gel loading on the basis of equal protein. PSII core (D1, D2) and antennae (Lhcb2) proteins were enriched in the grana core fraction, whereas PSI proteins (PsaB, PsaD, Lhca1), ATP synthase (AtpB) and cytochrome *b_6_f* (PetD) were more highly represented in nonappressed membranes (refined margins, curvature, and stromal lamellae), as expected. However, the considerable amount of PSI (PsaB, PsaD) and ATP synthase (AtpB) proteins in the grana core fraction confirmed this fraction to be contaminated with nonappressed membranes (Figure 1C). This contamination likely derived from incomplete solubilization of the peripheral regions of the grana due to the low detergent concentration needed to resolve the refined margin and curvature fractions.

The goal of this study was to examine the distribution of biogenesis factors among the different subdomains of nonappressed membranes, particularly the curvature and refined margin domains. Markers for these fractions partitioned as expected. CURT1A, a marker for the curvature domain (Armbruster et al., 2013), was strongly enriched in our curvature fraction (Figure 1C). Subunits of the cytochrome *b_6_f* and ATP synthase complexes were also enriched in the curvature fraction whereas PSI subunits were depleted from this fraction, as reported for the analogous fractions described previously (Trotta et al., 2019). VAR1/Ftsh5, a subunit of the protease that degrades photodamaged D1 (reviewed in Kato and Sakamoto, 2018), was enriched in the refined margin and stromal lamellae fractions, consistent with the partitioning of FtsH following protocols that produced a margin fraction that included the curvature domain (Puthiyaveetil et al., 2014; Nishioka et al., 2021). Most importantly, these results show that our fractionation successfully resolved the traditional margin fraction into two components: one of them enriched in FtsH and PSI (refined margins), the other enriched in CURT1, cyt *b_6_f*, and ATP synthase (curvature).

The assembly state of PSII changes in response to D1 photodamage to facilitate PSII repair (reviewed in Jarvi et al., 2015; Theis and Schroda, 2016). Two intermediates in PSII repair, the monomeric core complex and the CP43-free core, were highly enriched in a margin fraction that included the curvature domain (Koochak et al., 2019). The considerable representation of PSII core subunits D1 and D2 in our refined margin and/or curvature fractions (Figure 1C) is consistent with those results. Our data suggest that the abundance of D1 and D2 in the curvature fraction increases following illumination, although additional experimentation would be required to confirm this. Interestingly, two subunits of PSII’s "oxygen evolving complex" fractionated differently from one another: OE33 (also known as PsbO) partitioned similarly to D1 and D2, whereas OE23 (also known as PsbP) was enriched in the curvature fraction in both dark-adapted and illuminated plants. This finding, in conjunction with results presented below, adds to prior evidence that OE23/PsbP functions in the assembly of PSII independent of its role in mature PSII (Allahverdiyeva et al., 2013; Roose et al., 2016).

In summary, the partitioning of marker proteins among our curvature, refined margin, and stromal lamellae fractions as well as their chlorophyll *a/b* ratios are generally consistent with those reported previously for the corresponding fractions (Trotta et al., 2019). The small differences observed could result from our use of GDN instead of digitonin and plants at an earlier developmental stage. Our results confirm that the refined margin and curvature subfractions have distinct compositions, and support the view that the classic "margin" fraction includes both of these subfractions.

### Partitioning of Proteins Involved in Thylakoid Membrane Biogenesis Among Stroma-Exposed Thylakoid Subfractions

We next examined the localization of proteins involved in the biogenesis and repair of the photosynthetic apparatus, sampling proteins involved in protein synthesis, protein targeting, and the assembly of photosynthetic complexes (Figure 2). Plastid ribosomes synthesize core subunits of PSI, PSII, cyt *b_6_f* and ATP synthase, many of which co-translationally integrate into the thylakoid membrane. Ribosomes engaged in the synthesis of such proteins are bound to stroma-exposed membranes (Chua et al., 1973; Yamamoto et al., 1981; Zoschke and Barkan, 2015). Consistent with this, two plastid ribosomal proteins, RPS1 and RPS5, were found predominantly in our stromal lamellae fraction but with considerable representation in the refined margin fraction as well (Figure 2).

**Figure 2.**
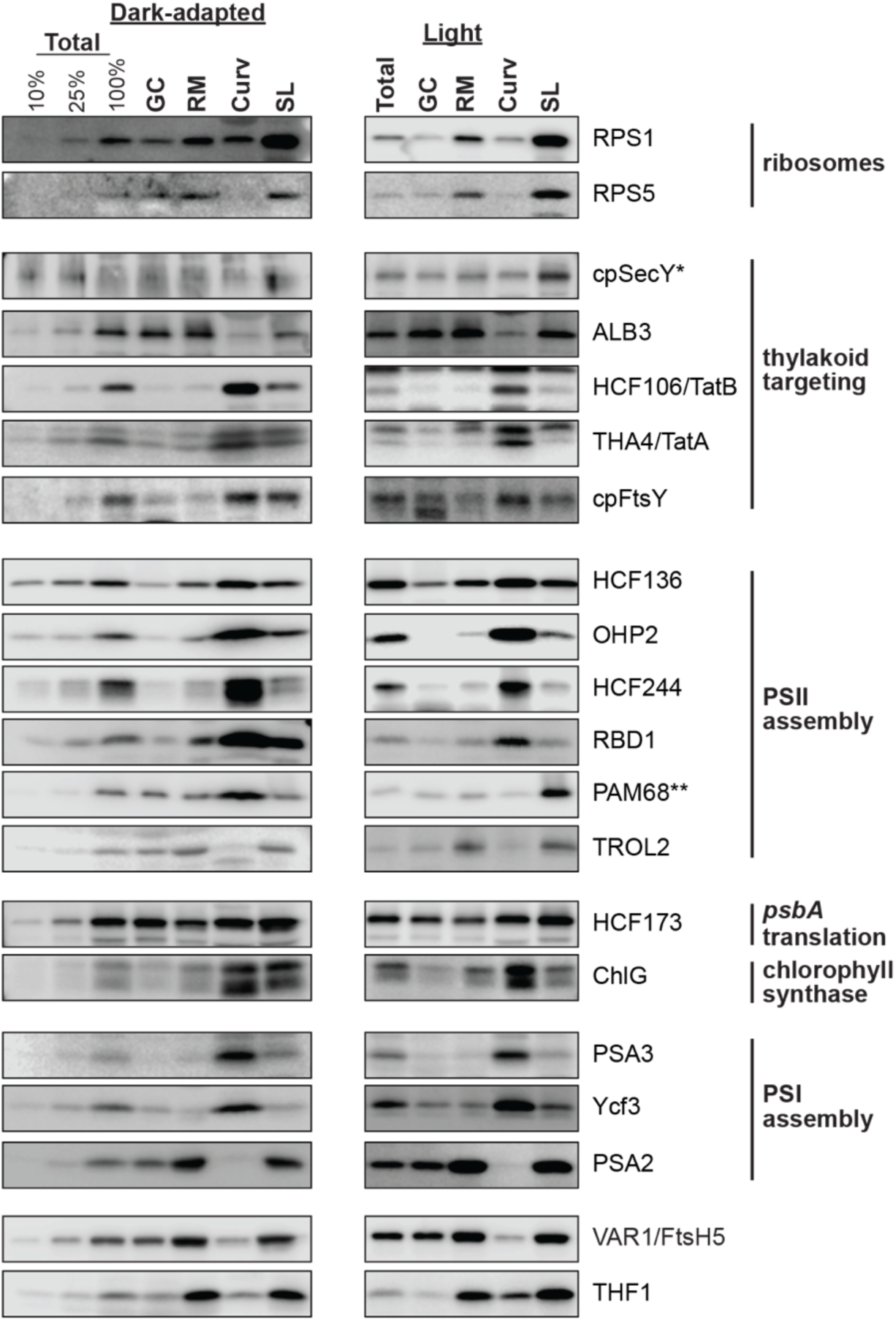
Immunoblot analysis of thylakoid biogenesis proteins in thylakoid subfractions. Replicates of the same samples described in Figure 1 were probed with antibodies to the indicated proteins. The VAR1/Ftsh5 data shown in Figure 1C is shown again here, to facilitate comparison with THF1 with whom it may interact. *SecY suffered degradation specifically in the fractions from dark-adapted plants. **PAM68 stood apart from the other proteins examined in that it fractionated in thylakoid membranes very differently in dark-adapted and illuminated plants.

Proteins are targeted to the thylakoid membrane via several mechanisms of cyanobacterial ancestry (reviewed in Celedon and Cline, 2013; Ziehe et al., 2018; Ries et al., 2020). We probed our fractions for components of each of these pathways. ALB3 and cpSecY, which function broadly in the co- and post-translational integration of proteins into the thylakoid membrane (Roy and Barkan, 1998; Ziehe et al., 2018; Ries et al., 2020; Ackermann et al., 2021), were distributed among thylakoid subfractions similarly to the plastid ribosomal proteins. This distribution is consistent with their known interactions with ribosomes (Ries et al., 2020; Ackermann et al., 2021) and with the prior localization of ALB3 to stromal lamellae and crude margins (Garcia-Cerdan et al., 2019). Curiously, cpSecY suffered degradation specifically in the membranes from dark-adapted plants, and was the only protein we examined to behave in that way. Given that plastid translation is globally repressed when plants are shifted to the dark (Chotewutmontri and Barkan, 2018), it seems plausible that decreased engagement of cpSecY with ribosomes during dark adaptation rendered cpSecY susceptible to proteolysis during the fractionation protocol.

The TAT translocon functions in the post-translational transport of specific folded proteins across the thylakoid membrane (Celedon and Cline, 2013). Two TAT subunits, HCF106 and THA4, were strongly enriched in the curvature subfraction (Figure 2), clearly separating from cpSecY, ALB3 and ribosomal proteins. Localization of TAT subunits to the curvature fraction raises the intriguing possibility that the unusual properties of the curvature domain are relevant to the membrane-destabilization mechanism for TAT transport (Asher and Theg, 2021; Hao et al., 2022). The SRP receptor cpFtsY was also enriched in the curvature fraction but with greater presence in margins and stromal lamellae than TAT subunits. The broader distribution of cpFtsY among the different nonappressed membrane fractions may reflect its participation in both co- and post-translational targeting of proteins to the thylakoid membrane (Asakura et al., 2004; Walter et al., 2015; Ziehe et al., 2018; Hristou et al., 2019).

Photosystems I and II are large macromolecular complexes whose assembly is facilitated by numerous accessory factors (Komenda and Sobotka, 2019; Nellaepalli et al., 2021; Komenda et al., 2024; Zhang et al., 2024). We probed our fractions for six proteins involved in PSII assembly (Figure 2). HCF136, which acts very early in the assembly of D1 into the PSII reaction center (Meurer et al., 1998; Plücken et al., 2002; Zhao et al., 2023), was distributed among the refined margin, curvature, and stromal lamellae fractions, consistent with the prior localization of HCF136 to crude margins and stromal lamellae (Garcia-Cerdan et al., 2019). HCF244, OHP2, and RBD1 also mediate early steps in assembly of the PSII reaction center and had previously been localized to crude margin fractions (Knoppova et al., 2014; Hey and Grimm, 2018; Garcia-Cerdan et al., 2019; Li et al., 2019; Maeda et al., 2022; Calderon et al., 2023). These three proteins localized almost exclusively to our curvature subfraction (Figure 2), contrasting with the broader distribution of HCF136 among nonappressed membrane fractions. Of possible relevance to this difference, the Synechocystis HCF136 ortholog (Ycf48) acts at multiple stages of PSII assembly, beginning with D1’s transit through the YidC insertase (ortholog of ALB3) (Yu et al., 2018) and extending past its association with the Ycf39 complex (ortholog of HCF244) (Komenda et al., 2024). That ALB3 was largely excluded from the curvature fraction, HCF244, OHP2, and RBD1 were largely excluded from refined margins and stromal lamellae, and HCF136 was distributed among all of these fractions is consistent with the multifarious roles for Ycf48/HCF136 discerned in Synechocystis.

PAM68 acts later in PSII assembly than HCF136, OHP2, HCF244, and RBD1, facilitating the incorporation of CP47 into the RC47 intermediate (Armbruster et al., 2010; Bučinská et al., 2018; Keller et al., 2023; Zhang et al., 2024). In dark-adapted plants, PAM68 was distributed among the three nonappressed fractions similarly to HCF136 (Figure 2). However, illumination caused PAM68 to shift almost entirely to stromal lamellae, a pattern that was strikingly different from the other PSII assembly factors we examined. The partitioning of PAM68 in illuminated plants was similar to that of ribosomal proteins, consistent with the fact that PAM68 copurifies with ribosomes (Bučinská et al., 2018; Zhang et al., 2024). Thus, the light-induced shift in PAM68’s localization may be linked to the global increase in chloroplast translation rates when plants are shifted from dark to light (Chotewutmontri and Barkan, 2018). A protein denoted TROL2, which has been proposed to act late in PSII assembly by promoting multimerization (Li et al., 2023), was found almost exclusively in the refined margin and stromal lamellae. The exclusion of TROL2 from the curvature fraction in both dark-adapted and illuminated plants set it apart from all other PSII assembly factors we examined.

We also examined the distribution of ChlG and HCF173 (Figure 2), whose functions have been tied to the HCF244 complex but that are not themselves assembly factors. ChlG, which catalyzes the last step in chlorophyll synthesis, copurified with the HCF244 ortholog in Synechocystis (Chidgey et al., 2014) and colocalized with the HCF244 complex in a crude margin fraction in Arabidopsis (Maeda et al., 2022). Like HCF244 and OHP2, ChlG was enriched in our curvature fraction. However, the partitioning of ChlG was most similar to that of HCF136, in that a considerable portion of both proteins was also found in the refined margins and stromal lamellae. HCF173 binds the chloroplast *psbA* mRNA and is required for its translation (Schult et al., 2007; McDermott et al., 2019). There is evidence that HCF173’s activity is regulated via an autoregulatory circuit centered in the HCF244/OHP2 complex (Chotewutmontri and Barkan, 2020; Rojas et al., 2024), but the mechanism underlying this crosstalk is unknown. Unlike the HCF244 complex which was found almost exclusively in the curvature fraction, HCF173 was distributed among all three nonappressed membrane fractions. This distribution is consistent with a transiting of HCF173-bound *psbA* polysomes from stromal lamellae (primary site of ribosomes) to the curvature fraction (primary site of HCF244 complex) during the course of D1 synthesis.

We next examined the partitioning of three PSI assembly factors (Figure 2). PSA3 and Ycf3, which act early in the assembly of PSI’s reaction center (Shen et al., 2017; Nellaepalli et al., 2018), were found primarily in the curvature fraction. By contrast, PSA2, which facilitates a late step in PSI assembly (Fristedt et al., 2014), was excluded from the curvature fraction and abundant in refined margins and stromal lamellae. In fact, the distribution of PSA2 was similar to that of PSI subunits (see PsaD and Lhca1 in Fig. 1), consistent with a function associated with mature or nearly mature PSI.

Finally, we addressed the localization of THF1, a membrane-extrinsic thylakoid protein whose loss causes leaf variegation and disruptions in thylakoid biogenesis (Wang et al., 2004). The cyanobacterial THF1 ortholog associates with the FtsH protease and is required for normal FtsH accumulation (Beckova et al., 2017). A similar role in chloroplasts is suggested by the fact that Arabidopsis *thf1* and *ftsH* mutants exhibit similar variegation phenotypes (Chen et al., 2000; Sakamoto et al., 2003). THF1 was distributed among the three nonappressed membrane fractions in a pattern that resembled that of the FtsH subunit VAR1/FtsH5 (Figure 2), consistent with the possibility that their functions are related. The partitioning of THF1 also resembled that of the ALB3 insertase, but differed from all other biogenesis factors we examined.

## Discussion

Results presented here elucidate the spatial organization of processes underlying the assembly and repair of the photosynthetic apparatus in plant chloroplasts. We focused on the partitioning of biogenesis factors among the refined margin and curvature fractions, recognizing that the morphological features represented in these fractions remain incompletely defined. For example, the material that copurifies with CURT1 in the "curvature" fraction might be localized either to the highly curved grana periphery or to a distinct thylakoid subdomain with similar solubilization and sedimentation characteristics. This caveat notwithstanding, the various fractionation patterns we observed highlight the spatial heterogeneity of biogenesis and repair processes in the thylakoid membrane. New insights from our survey are summarized below.

The partitioning of structural components of the photosynthetic apparatus among our fractions was similar to that reported previously for dark-adapted plants (Trotta et al., 2019): a subunit of the cyt *b_6_f* complex was concentrated in the curvature fraction and was absent from refined margins, whereas subunits of PSI were absent from the curvature fraction and concentrated in margins. We found, in addition, that ATP synthase subunits were concentrated in the curvature fraction and that none of these patterns changed substantively upon illumination. PSI, cyt *b_6_f* and ATP synthase, in addition to residing in stromal lamellae, have been reported to colocalize in margins (Koochak et al., 2019; Rantala et al., 2020). However, our resolution of the margin fraction into two components shows that PSI is largely segregated from ATP synthase and cyt *b_6_f* in membrane fractions with distinct properties. The enrichment of the PSII subunit OE23/PsbP in the curvature fraction set it apart from its OE33 partner and from other PSII subunits, supporting the previous conjecture that OE23/PsbP assists in PSII assembly (Allahverdiyeva et al., 2013). Several OE23/PsbP homologs have known functions in photosystem assembly (Liu et al., 2012; Knoppova et al., 2016; Roose et al., 2016; Che et al., 2020), so an analogous role for OE23/PsbP would not be surprising.

The chloroplast FtsH protease plays a central role in PSII repair by degrading photodamaged D1 (reviewed in Kato and Sakamoto, 2018). Its VAR1/FtsH5 subunit was enriched in the refined margin and stromal lamellae fractions, consistent with the prior localization of FtsH to a traditional margin fraction and stromal lamellae (Puthiyaveetil et al., 2014; Nishioka et al., 2021). However, VAR1/FtsH5 fractionated similarly in dark-adapted and illuminated plants, contrasting with a report that FtsH partially redistributes from stromal lamellae to margins when dark-adapted plants are shifted to high intensity light (Puthiyaveetil et al., 2014). Our use of a moderate light intensity that is standard for Arabidopsis growth may account for this difference. In fact, the only protein in our survey that unambiguously changed its distribution in response to illumination was the PSII assembly factor PAM68: in illuminated plants PAM68 cofractionated with ribosomal proteins, consistent with its association with ribosomes (Bučinská et al., 2018; Zhang et al., 2024), whereas in dark-adapted plants PAM68 was enriched in the curvature fraction. The mechanism underlying this behavior and its functional significance would be interesting to explore in the future.

After FtsH degrades photodamaged D1, newly synthesized D1 is assembled into a D1-less PSII repair intermediate in a process believed to involve the same factors that promote early steps of *de novo* PSII assembly, including HCF136, HCF244, OHP2, and RBD1 (Komenda et al., 2024). It is interesting in this context that these factors partitioned among thylakoid subfractions in different ways: VAR1/FtsH5 localized to refined margins and stromal lamellae, HCF244, OHP2, and RBD1 were largely restricted to the curvature fraction, and HCF136 was found in all three fractions. These fractionation patterns lend credence to the suggestion (Puthiyaveetil et al., 2014) that consecutive steps in PSII repair are physically segregated in a manner that reduces opportunities for wasteful back reactions. This correlation extends to ribosomal proteins and the ALB3 insertase, involved in the earliest steps of photosystem synthesis and assembly, which were depleted from the curvature fraction. HCF173, the translational activator that regulates D1 synthesis in response to light-induced D1 damage (Schult et al., 2007; Rojas et al., 2024), was well represented in all three nonappressed membrane fractions, consistent with its presence on *psbA* mRNA undergoing translation and its proposed regulation by the HCF244/OHP2 complex (McDermott et al., 2019; Chotewutmontri and Barkan, 2020).

Our results suggest a similar spatial segregation of different steps in the assembly of PSI. PSA3 and YCF3, involved in early steps in PSI assembly (Shen et al., 2017; Nellaepalli et al., 2018), were enriched in the curvature fraction whereas PSA2, which acts late in PSI assembly (Fristedt et al., 2014), was absent from the curvature fraction and distributed between refined margins and stromal lamellae. It would be informative to examine the distribution of additional PSI assembly factors whose order of action has been reported (e.g. Nellaepalli et al., 2021; Rolo et al., 2024; Zhang et al., 2024).

In summary, the results reported here offer insights into the physical arrangement of photosystem biogenesis and repair processes in the thylakoid membrane system of plant chloroplasts. Our results highlight the curvature fraction as having special pertinence to early events in photosystem assembly, and suggest that different assembly steps are physically segregated in a manner that reinforces the vectorial nature of assembly pathways. In the green alga *Chlamydomonas reinhardtii*, whose thylakoid membrane architecture is quite different than that in plants, fluorescence microscopy has localized photosystem biogenesis to a specialized region (Schottkowski et al., 2012; Sun et al., 2019). Our results suggest that the curvature fraction of plant thylakoid membranes may occupy an analogous functional niche. High resolution *in situ* imaging approaches will be needed to clarify the relationships among the morphological features of plant thylakoid membranes and the biochemically-defined stromal lamellae, refined margin, and curvature fractions. Furthermore, elucidating the mechanisms underlying the diverse partitioning patterns of biogenesis factors among different nonappressed membrane fractions will be essential for understanding the orchestration of thylakoid biogenesis.

## Accession Numbers

ALB3 AT2G28800

ChlG AT3G51820

cpFtsY AT2G45770

cpSecY AT2G18710

HCF106 AT5G52440

HCF136 AT5G23120

HCF173 AT1G16720

HCF244 AT4G35250

OHP2 AT1G34000

PAM68 AT4G19100

PSA2 AT2G34860

PSA3 AT3G55250

RBD1 AT1G54500

THA4 AT5G28750

THF1 AT2G20890

TROL2 AT3G25480

YCF3 ATCG00360

## Acknowledgements

We are grateful to Rosalind Williams-Carrier (University of Oregon) for expert technical assistance, and to the scientists who generously provided antibodies we used in this study: Carole Dabney-Smith (Miami University), Misty Moore and Ralph Henry (University of Arkansas), Wataru Sakamoto (Okayama University), Peter Weshoff (University of Düsseldorf), Kris Niyogi (University of California). This work was supported by grants to A.B. from the Department of Energy (DE-SC0018916) and National Science Foundation (MCB-2034758).

## Author Contributions

PC designed the research, performed the research, analyzed the data, and edited the paper. AB designed the research, analyzed the data and wrote the paper.

## Conflict of Interest

The authors have no conflicts of interest to declare.

## Notes

### Competing Interest Statement

The authors have declared no competing interest.

## Literature Cited

Ackermann B, Dunschede B, Pietzenuk B, Justesen BH, Krämer U, Hofmann E, Pomorski TG, Schünemann D (2021) Chloroplast Ribosomes Interact With the Insertase Alb3 in the Thylakoid Membrane. Front Plant Sci 12, 781857

Allahverdiyeva Y, Suorsa M, Rossi F, Pavesi A, Kater MM, Antonacci A, Tadini L, Pribil M, Schneider A, Wanner G, Leister D, Aro EM, Barbato R, Pesaresi P (2013) Arabidopsis plants lacking PsbQ and PsbR subunits of the oxygen-evolving complex show altered PSII super-complex organization and short-term adaptive mechanisms. Plant J 75, 671–684

Anderson JM, Andersson B (1988) The dynamic photosynthetic membrane and regulation of solar energy conversion. Trends Biochem Sci 13, 351–355

Armbruster U, Labs M, Pribil M, Viola S, Xu W, Scharfenberg M, Hertle AP, Rojahn U, Jensen PE, Rappaport F, Joliot P, Dormann P, Wanner G, Leister D (2013) Arabidopsis CURVATURE THYLAKOID1 proteins modify thylakoid architecture by inducing membrane curvature. Plant Cell 25, 2661–2678

Armbruster U, Zuhlke J, Rengstl B, Kreller R, Makarenko E, Ruhle T, Schunemann D, Jahns P, Weisshaar B, Nickelsen J, Leister D (2010) The Arabidopsis thylakoid protein PAM68 is required for efficient D1 biogenesis and photosystem II assembly. Plant Cell 22, 3439–3460

Asakura Y, Hirohashi T, Kikuchi S, Belcher S, Osborne E, Yano S, Terashima I, Barkan A, Nakai M (2004) Maize mutants lacking chloroplast FtsY exhibit pleiotropic defects in the biogenesis of thylakoid membranes. Plant Cell 16, 201–214

Asher AH, Theg SM (2021) Electrochromic shift supports the membrane destabilization model of Tat-mediated transport and shows ion leakage during Sec transport. Proc Natl Acad Sci U S A 118

Austin JR, 2nd, Staehelin LA (2011) Three-dimensional architecture of grana and stroma thylakoids of higher plants as determined by electron tomography. Plant Physiol 155, 1601-1611

Barkan A (1998) Approaches to investigating nuclear genes that function in chloroplast biogenesis in land plants. Methods Enzymol 297, 38–57

Beckova M, Yu J, Krynicka V, Kozlo A, Shao S, Konik P, Komenda J, Murray JW, Nixon PJ (2017) Structure of Psb29/Thf1 and its association with the FtsH protease complex involved in photosystem II repair in cyanobacteria. Philos Trans R Soc Lond B Biol Sci 372

Bučinská L, Kiss E, Koník P, Knoppová J, Komenda J, Sobotka R (2018) The Ribosome-Bound Protein Pam68 Promotes Insertion of Chlorophyll into the CP47 Subunit of Photosystem II. Plant Physiol 176, 2931–2942

Calderon RH, de Vitry C, Wollman FA, Niyogi KK (2023) Rubredoxin 1 promotes the proper folding of D1 and is not required for heme b(559) assembly in Chlamydomonas photosystem II. J Biol Chem 299, 102968

Calderon RH, Garcia-Cerdan JG, Malnoe A, Cook R, Russell JJ, Gaw C, Dent RM, de Vitry C, Niyogi KK (2013) A conserved rubredoxin is necessary for photosystem II accumulation in diverse oxygenic photoautotrophs. J Biol Chem 288, 26688–26696

Celedon JM, Cline K (2013) Intra-plastid protein trafficking: how plant cells adapted prokaryotic mechanisms to the eukaryotic condition. Biochim Biophys Acta 1833, 341–351

Che Y, Kusama S, Matsui S, Suorsa M, Nakano T, Aro EM, Ifuku K (2020) Arabidopsis PsbP-Like Protein 1 Facilitates the Assembly of the Photosystem II Supercomplexes and Optimizes Plant Fitness under Fluctuating Light. Plant Cell Physiol 61, 1168–1180

Chen M, Choi Y, Voytas DF, Rodermel S (2000) Mutations in the Arabidopsis VAR2 locus cause leaf variegation due to the loss of a chloroplast FtsH protease. Plant J 22, 303–313

Chidgey JW, Linhartova M, Komenda J, Jackson PJ, Dickman MJ, Canniffe DP, Konik P, Pilny J, Hunter CN, Sobotka R (2014) A cyanobacterial chlorophyll synthase-HliD complex associates with the Ycf39 protein and the YidC/Alb3 insertase. Plant Cell 26, 1267–1279

Chotewutmontri P, Barkan A (2018) Multilevel effects of light on ribosome dynamics in chloroplasts program genome-wide and psbA-specific changes in translation. PLoS Genet 14, e1007555

Chotewutmontri P, Barkan A (2020) Light-induced *psbA* translation in plants is triggered by photosystem II damage via an assembly-linked autoregulatory circuit. Proc Natl Acad Sci U S A 117, 21775–21784

Chotewutmontri P, Williams-Carrier R, Barkan A (2020) Exploring the link between photosystem II assembly and translation of the chloroplast *psbA* mRNA. Plants 9, pii: E152

Chua NH, Blobel G, Siekevitz P, Palade GE (1973) Attachment of chloroplast polysomes to thylakoid membranes in Chlamydomonas reinhardtii. Proc Natl Acad Sci U S A 70, 1554–1558

Daum B, Kuhlbrandt W (2011) Electron tomography of plant thylakoid membranes. J Exp Bot 62, 2393–2402

Flannery SE, Hepworth C, Wood WHJ, Pastorelli F, Hunter CN, Dickman MJ, Jackson PJ, Johnson MP (2021) Developmental acclimation of the thylakoid proteome to light intensity in Arabidopsis. Plant J 105, 223–244

Fristedt R, Williams-Carrier R, Merchant SS, Barkan A (2014) A thylakoid membrane protein harboring a DnaJ-type zinc finger domain is required for photosystem I accumulation in plants. J Biol Chem 289, 30657–30667

Garcia-Cerdan JG, Furst AL, McDonald KL, Schunemann D, Francis MB, Niyogi KK (2019) A thylakoid membrane-bound and redox-active rubredoxin (RBD1) functions in de novo assembly and repair of photosystem II. Proc Natl Acad Sci U S A 116, 16631–16640

Hao B, Zhou W, Theg SM (2022) Hydrophobic mismatch is a key factor in protein transport across lipid bilayer membranes via the Tat pathway. J Biol Chem 298, 101991

Hey D, Grimm B (2018) ONE-HELIX PROTEIN2 (OHP2) Is Required for the Stability of OHP1 and Assembly Factor HCF244 and Is Functionally Linked to PSII Biogenesis. Plant Physiol 177, 1453–1472

Höhner R, Pribil M, Herbstova M, Lopez LS, Kunz HH, Li M, Wood M, Svoboda V, Puthiyaveetil S, Leister D, Kirchhoff H (2020) Plastocyanin is the long-range electron carrier between photosystem II and photosystem I in plants. Proc Natl Acad Sci U S A 117, 15354–15362

Hristou A, Gerlach I, Stolle DS, Neumann J, Bischoff A, Dünschede B, Nowaczyk MM, Zoschke R, Schünemann D (2019) Ribosome-Associated Chloroplast SRP54 Enables Efficient Cotranslational Membrane Insertion of Key Photosynthetic Proteins. Plant Cell 31, 2734–2750

Jarvi S, Suorsa M, Aro EM (2015) Photosystem II repair in plant chloroplasts--Regulation, assisting proteins and shared components with photosystem II biogenesis. Biochim Biophys Acta 1847, 900–909

Kato Y, Sakamoto W (2018) FtsH Protease in the Thylakoid Membrane: Physiological Functions and the Regulation of Protease Activity. Front Plant Sci 9, 855

Keller JM, Frieboes MJ, Jodecke L, Kappel S, Wulff N, Rindfleisch T, Sandoval-Ibanez O, Gerlach I, Thiele W, Bock R, Eirich J, Finkemeier I, Schunemann D, Zoschke R, Schottler MA, Armbruster U (2023) Eukaryote-specific assembly factor DEAP2 mediates an early step of photosystem II assembly in Arabidopsis. Plant Physiol 193, 1970–1986

Kirchhoff H (2019) Chloroplast ultrastructure in plants. New Phytol 223, 565–574

Kirchhoff H, Hall C, Wood M, Herbstova M, Tsabari O, Nevo R, Charuvi D, Shimoni E, Reich Z (2011) Dynamic control of protein diffusion within the granal thylakoid lumen. Proc Natl Acad Sci U S A 108, 20248–20253

Kirchhoff H, Li M, Puthiyaveetil S (2017) Sublocalization of Cytochrome b(6)f Complexes in Photosynthetic Membranes. Trends Plant Sci 22, 574–582

Knoppova J, Sobotka R, Tichy M, Yu J, Konik P, Halada P, Nixon PJ, Komenda J (2014) Discovery of a chlorophyll binding protein complex involved in the early steps of photosystem II assembly in Synechocystis. Plant Cell 26, 1200–1212

Knoppova J, Yu J, Konik P, Nixon PJ, Komenda J (2016) CyanoP is Involved in the Early Steps of Photosystem II Assembly in the Cyanobacterium Synechocystis sp. PCC 6803. Plant Cell Physiol 57, 1921-1931

Komenda J, Sobotka R (2019) Chlorophyll-binding subunits of photosystem I and II: Biosynthesis, chlorophyll incorporation and assembly. In Advances in Botanical Res. Academic Press

Komenda J, Sobotka R, Nixon PJ (2024) The biogenesis and maintenance of photosystem II: recent advances and current challenges. Plant Cell

Koochak H, Puthiyaveetil S, Mullendore DL, Li M, Kirchhoff H (2019) The structural and functional domains of plant thylakoid membranes. Plant J 97, 412–429

Lee HJ, Lee HS, Youn T, Byrne B, Chae PS (2022) Impact of novel detergents on membrane protein studies. Chem 8, 980–1013

Li M, Mukhopadhyay R, Svoboda V, Oung HMO, Mullendore DL, Kirchhoff H (2020) Measuring the dynamic response of the thylakoid architecture in plant leaves by electron microscopy. Plant Direct 4, e00280

Li W, Guo J, Han X, Da X, Wang K, Zhao H, Huang ST, Li B, He H, Jiang R, Zhou S, Yan P, Chen T, He Y, Xu J, Liu Y, Wu Y, Shou H, Wu Z, Mao C, Mo X (2023) A novel protein domain is important for photosystem II complex assembly and photoautotrophic growth in angiosperms. Mol Plant 16, 374–392

Li Y, Liu B, Zhang J, Kong F, Zhang L, Meng H, Li W, Rochaix JD, Li D, Peng L (2019) OHP1, OHP2, and HCF244 Form a Transient Functional Complex with the Photosystem II Reaction Center. Plant Physiol 179, 195-208

Liu J, Yang H, Lu Q, Wen X, Chen F, Peng L, Zhang L, Lu C (2012) PsbP-domain protein1, a nuclear-encoded thylakoid lumenal protein, is essential for photosystem I assembly in Arabidopsis. Plant Cell 24, 4992–5006

Maeda H, Takahashi K, Ueno Y, Sakata K, Yokoyama A, Yarimizu K, Myouga F, Shinozaki K, Ozawa SI, Takahashi Y, Tanaka A, Ito H, Akimoto S, Takabayashi A, Tanaka R (2022) Characterization of photosystem II assembly complexes containing ONE-HELIX PROTEIN1 in Arabidopsis thaliana. J Plant Res 135, 361–376

McDermott JJ, Watkins KP, Williams-Carrier R, Barkan A (2019) Ribonucleoprotein capture by in vivo expression of a designer pentatricopeptide repeat protein in Arabidopsis. Plant Cell 31, 1723–1733

Meurer J, Plücken H, Kowallik K, Westhoff P (1998) A nuclear-encoded protein of prokaryotic origin is essential for the stability of photosystem II in Arabidopsis thaliana. EMBO J 17, 5286–5297

Moore M, Harrison MS, Peterson EC, Henry R (2000) Chloroplast Oxa1p homolog albino3 is required for post-translational integration of the light harvesting chlorophyll-binding protein into thylakoid membranes. J Biol Chem 275, 1529–1532

Mori H, Summer EJ, Ma X, Cline K (1999) Component specificity for the thylakoidal Sec and Delta pH-dependent protein transport pathways. J Cell Biol 146, 45–56

Nellaepalli S, Kim RG, Grossman AR, Takahashi Y (2021) Interplay of four auxiliary factors is required for the assembly of photosystem I reaction center subcomplex. Plant J 106, 1075–1086

Nellaepalli S, Ozawa SI, Kuroda H, Takahashi Y (2018) The photosystem I assembly apparatus consisting of Ycf3-Y3IP1 and Ycf4 modules. Nat Commun 9, 2439

Nishimura K, Kato Y, Sakamoto W (2016) Chloroplast Proteases: Updates on Proteolysis within and across Suborganellar Compartments. Plant Physiol 171, 2280–2293

Nishioka K, Kato Y, Ozawa SI, Takahashi Y, Sakamoto W (2021) Phos-tag-based approach to study protein phosphorylation in the thylakoid membrane. Photosynth Res 147, 107–124

Ogawa Y, Iwano M, Shikanai T, Sakamoto W (2023) FZL, a dynamin-like protein localized to curved grana edges, is required for efficient photosynthetic electron transfer in Arabidopsis. Front Plant Sci 14, 1279699

Peltier JB, Ytterberg AJ, Sun Q, van Wijk KJ (2004) New functions of the thylakoid membrane proteome of Arabidopsis thaliana revealed by a simple, fast, and versatile fractionation strategy. J Biol Chem 279, 49367–49383

Plücken H, Müller B, Grohmann D, Westhoff P, Eichacker LA (2002) The HCF136 protein is essential for assembly of the photosystem II reaction center in Arabidopsis thaliana. FEBS Lett 532, 85–90

Porra RJ, Thompson WA, Kriedemann PE (1989) Determination of accurate extinction coefficients and simultaneous equations for assaying chlorophylls a and b extracted with four different solvents. BBA Bioenergetics 975, 384–394

Puthiyaveetil S, Tsabari O, Lowry T, Lenhert S, Lewis RR, Reich Z, Kirchhoff H (2014) Compartmentalization of the protein repair machinery in photosynthetic membranes. Proc Natl Acad Sci U S A 111, 15839–15844

Rantala M, Rantala S, Aro EM (2020) Composition, phosphorylation and dynamic organization of photosynthetic protein complexes in plant thylakoid membrane. Photochem Photobiol Sci 19, 604–619

Ries F, Herkt C, Willmund F (2020) Co-Translational Protein Folding and Sorting in Chloroplasts. Plants (Basel) 9

Rojas M, Chotewutmontri P, Barkan A (2024) Translational activation by a synthetic PPR protein elucidates control of psbA translation in Arabidopsis chloroplasts. Plant Cell

Rolo D, Sandoval-Ibanez O, Thiele W, Schottler MA, Gerlach I, Zoschke R, Schwartzmann J, Meyer EH, Bock R (2024) CO-EXPRESSED WITH PSI ASSEMBLY1 (CEPA1) is a photosystem I assembly factor in Arabidopsis. Plant Cell

Roose JL, Frankel LK, Mummadisetti MP, Bricker TM (2016) The extrinsic proteins of photosystem II: update. Planta 243, 889–908

Roy LM, Barkan A (1998) A secY homologue is required for the elaboration of the chloroplast thylakoid membrane and for normal chloroplast gene expression. J. Cell Biol. 141, 385–395

Ruban AV, Johnson MP (2015) Visualizing the dynamic structure of the plant photosynthetic membrane. Nat Plants 1, 15161

Sakamoto W, Zaltsman A, Adam Z, Takahashi Y (2003) Coordinated regulation and complex formation of yellow variegated1 and yellow variegated2, chloroplastic FtsH metalloproteases involved in the repair cycle of photosystem II in Arabidopsis thylakoid membranes. Plant Cell 15, 2843–2855

Schottkowski M, Peters M, Zhan Y, Rifai O, Zhang Y, Zerges W (2012) Biogenic membranes of the chloroplast in Chlamydomonas reinhardtii. Proc Natl Acad Sci U S A 109, 19286–19291

Schult K, Meierhoff K, Paradies S, Toller T, Wolff P, Westhoff P (2007) The nuclear-encoded factor HCF173 is involved in the initiation of translation of the *psbA* mRNA in Arabidopsis thaliana. Plant Cell 19, 1329–1346

Shen J, Williams-Carrier R, Barkan A (2017) PSA3, a Protein on the Stromal Face of the Thylakoid Membrane, Promotes Photosystem I Accumulation in Cooperation with the Assembly Factor PYG7. Plant Physiol 174, 1850-1862

Sun Y, Valente-Paterno MI, Bakhtiari S, Law C, Zhan Y, Zerges W (2019) Photosystem Biogenesis Is Localized to the Translation Zone in the Chloroplast of Chlamydomonas. Plant Cell

Theis J, Schroda M (2016) Revisiting the photosystem II repair cycle. Plant Signal Behav 11, e1218587

Tomizioli M, Lazar C, Brugiere S, Burger T, Salvi D, Gatto L, Moyet L, Breckels LM, Hesse AM, Lilley KS, Seigneurin-Berny D, Finazzi G, Rolland N, Ferro M (2014) Deciphering thylakoid sub-compartments using a mass spectrometry-based approach. Mol Cell Proteomics 13, 2147–2167

Trotta A, Bajwa AA, Mancini I, Paakkarinen V, Pribil M, Aro EM (2019) The Role of Phosphorylation Dynamics of CURVATURE THYLAKOID 1B in Plant Thylakoid Membranes. Plant Physiol 181, 1615–1631

Walker M, Roy L, Coleman E, Voelker R, Barkan A (1999) The maize *tha4* gene functions in sec-independent protein transport in chloroplasts and is related to *hcf106, tatA*, and *tatB*. J. Cell Biol. 147, 267–275

Walter B, Pieta T, Schünemann D (2015) Arabidopsis thaliana mutants lacking cpFtsY or cpSRP54 exhibit different defects in photosystem II repair. Front Plant Sci 6, 250

Wang Q, Sullivan RW, Kight A, Henry RL, Huang J, Jones AM, Korth KL (2004) Deletion of the chloroplast-localized Thylakoid formation1 gene product in Arabidopsis leads to deficient thylakoid formation and variegated leaves. Plant Physiol 136, 3594–3604

Yamamoto T, Burke J, Autz G, Jagendorf AT (1981) Bound Ribosomes of Pea Chloroplast Thylakoid Membranes: Location and Release in Vitro by High Salt, Puromycin, and RNase. Plant Physiol 67, 940–949

Yoshioka-Nishimura M (2016) Close Relationships Between the PSII Repair Cycle and Thylakoid Membrane Dynamics. Plant Cell Physiol 57, 1115–1122

Yoshioka-Nishimura M, Nanba D, Takaki T, Ohba C, Tsumura N, Morita N, Sakamoto H, Murata K, Yamamoto Y (2014) Quality control of photosystem II: direct imaging of the changes in the thylakoid structure and distribution of FtsH proteases in spinach chloroplasts under light stress. Plant Cell Physiol 55, 1255–1265

Yu J, Knoppova J, Michoux F, Bialek W, Cota E, Shukla MK, Straskova A, Pascual Aznar G, Sobotka R, Komenda J, Murray JW, Nixon PJ (2018) Ycf48 involved in the biogenesis of the oxygen-evolving photosystem II complex is a seven-bladed beta-propeller protein. Proc Natl Acad Sci U S A 115, E7824–E7833

Zhang A, Tian L, Zhu T, Li M, Sun M, Fang Y, Zhang Y, Lu C (2024) Uncovering the photosystem I assembly pathway in land plants. Nat Plants 10, 645–660

Zhang L, Ruan J, Gao F, Xin Q, Che LP, Cai L, Liu Z, Kong M, Rochaix JD, Mi H, Peng L (2024) Thylakoid protein FPB1 synergistically cooperates with PAM68 to promote CP47 biogenesis and Photosystem II assembly. Nat Commun 15, 3122

Zhao Z, Vercellino I, Knoppova J, Sobotka R, Murray JW, Nixon PJ, Sazanov LA, Komenda J (2023) The Ycf48 accessory factor occupies the site of the oxygen-evolving manganese cluster during photosystem II biogenesis. Nat Commun 14, 4681

Ziehe D, Dünschede B, Schünemann D (2018) Molecular mechanism of SRP-dependent light-harvesting protein transport to the thylakoid membrane in plants. Photosynth Res 138, 303–313

Zoschke R, Barkan A (2015) Genome-wide analysis of thylakoid-bound ribosomes in maize reveals principles of cotranslational targeting to the thylakoid membrane. Proc Natl Acad Sci U S A 112, E1678–1687

